# Silencing epigenetic writer DOT1L attenuates intimal hyperplasia

**DOI:** 10.1101/730572

**Authors:** Yitao Huang, Go Urabe, Mengxue Zhang, Jing Li, Bowen Wang, K. Craig Kent, Lian-Wang Guo

## Abstract

**Objective:** Histone methyltransferases are emerging targets for epigenetic therapy. DOT1L (disruptor of telomeric silencing 1-like) is the H3K79 methylation writer. We investigated its role in the development of intimal hyperplasia (IH).

**Approach and Results:** IH was induced via balloon angioplasty in rat carotid arteries. DOT1L and its catalytic products H3K79me2 and H3K79me3 (immunostaining) increased by 4.69 ±0.34, 2.38 ±0.052, and 3.07 ±0.27 fold, respectively, in injured (*versus* uninjured) carotid arteries at post-injury day 7. DOT1L silencing via shRNA-lentivirus infusion in injured arteries reduced DOT1L, H3K79me2, and IH at day 14 by 54.5%, 37.1%, and 76.5%, respectively. Moreover, perivascular administration of a DOT1L-selective inhibitor (EPZ5676) reduced H3K79me2, H3K79me3, and IH by 56.1%, 58.6%, and 39.9%, respectively.

**Conclusions:** DOT1L inhibition mitigates the development of IH.

## Introduction

(Neo)intimal hyperplasia (IH), which narrows vasculature lumen and obstructs blood flow, occurs in major cardiovascular diseases^1^. While epigenetic therapy emerges as new medicine^2^, epigenetic targets in vascular diseases are under-exploited.

DOT1L catalyzes methylation at H3K79. Unlike other histone methylation sites located on the unstructured histone tail, H3K79 methylation occurs structure-dependently on the H3 globular domain^3^, and its erasers and readers (if any) remain unidentified^4^. Whether DOT1L plays a role in IH was not known. There is a strong correlation between H3K79 methylation and cancer progression^4^. We hypothesized that DOT1L plays a role in neointima formation.

In this study, we detected elevated DOT1L protein and function (typically measured as H3K79 dimethylation)^5^ in rat and human pathological arteries. Silencing DOT1L diminished IH in the model of rat carotid artery injury. We were also able to reduce IH by pharmacologically inhibiting DOT1L. These results support an important role of DOT1L in IH.

## Methods

Sprague-Dawley rats underwent balloon injury in the common carotid artery. To silence DOT1L in vivo, shRNA-expressing lentiviruses were infused into the endothelial-denuded artery wall, as we reported previously^6^. To pharmacologically inhibit the DOT1L methyltransferase activity, EPZ5676^7^ carried in thermo-sensitive hydrogel was applied around the injured artery^1^. For morphometric analysis, injured and uninjured arteries were collected at post-injury day 3, day 7, and day 14. Immunostaining and IH measurement (intima/media area ratio) were performed on paraffin sections^1^. One-way ANOVA (and Tukey’s post-hoc analysis) or unpaired Student t-test (and also non-parametric test) was applied for multi-group and two-group statistical analyses, respectively, as specified in each figure legend. More detailed methods can be found in Supplemental Materials.

## Results

To investigate a possible role of DOT1L in neointimal development, we used a well-established rat model of IH in which balloon angioplasty injures the carotid artery inducing neointima formation^1^. While neointima typically initiates at post-injury day 3, accelerates around day 7, and maximizes at day 14^1, 6^, we observed that immunostaining of DOT1L (Figure 1A, Figure S1) dramatically increased at day 3 (3.53 ±0.36 fold), peaked at day 7 (4.69 ±0.34 fold), and remained high at day 14 (3.62 ±0.36 fold), in injured arteries *vs* uninjured control.

**Figure 1.**
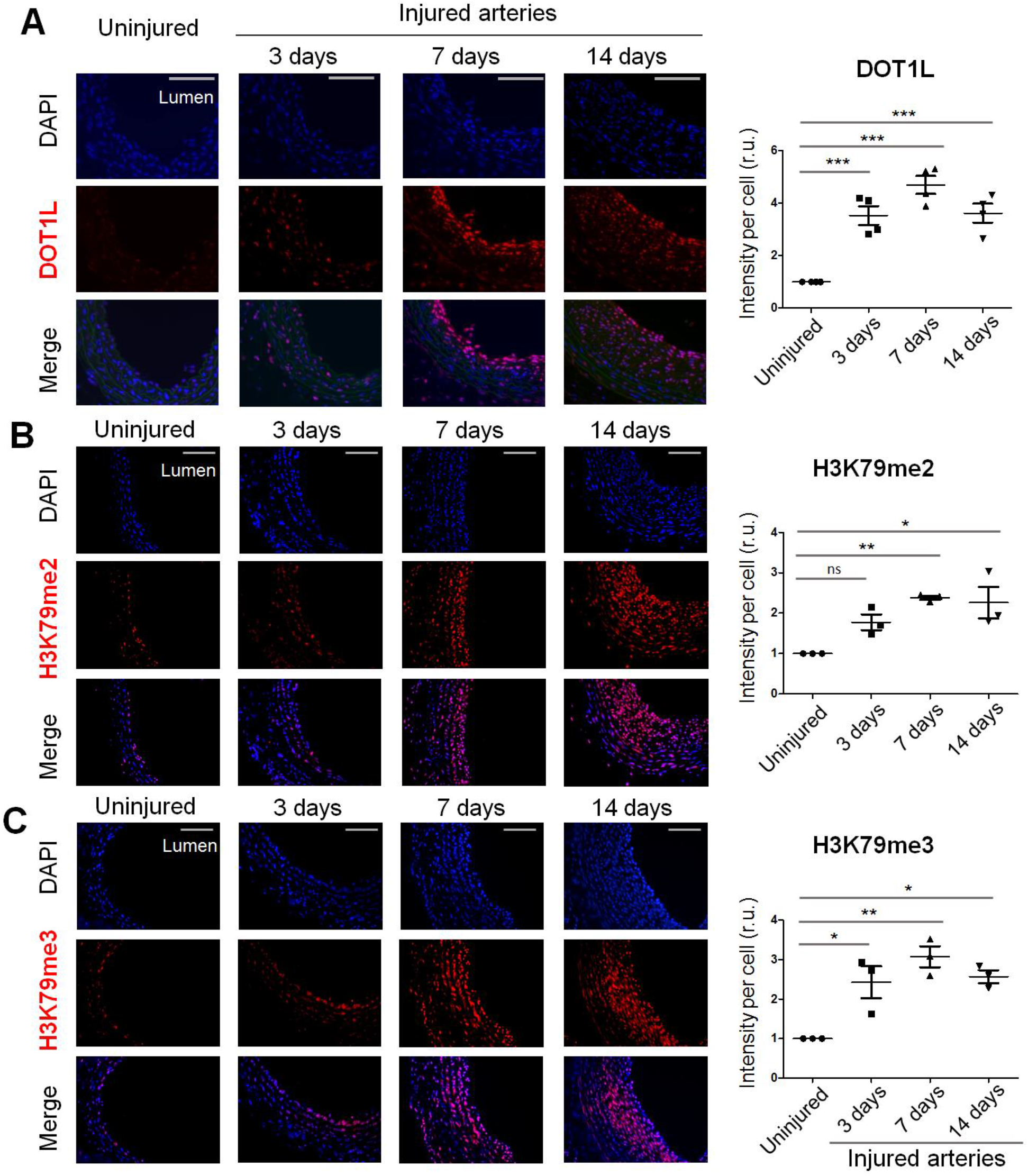
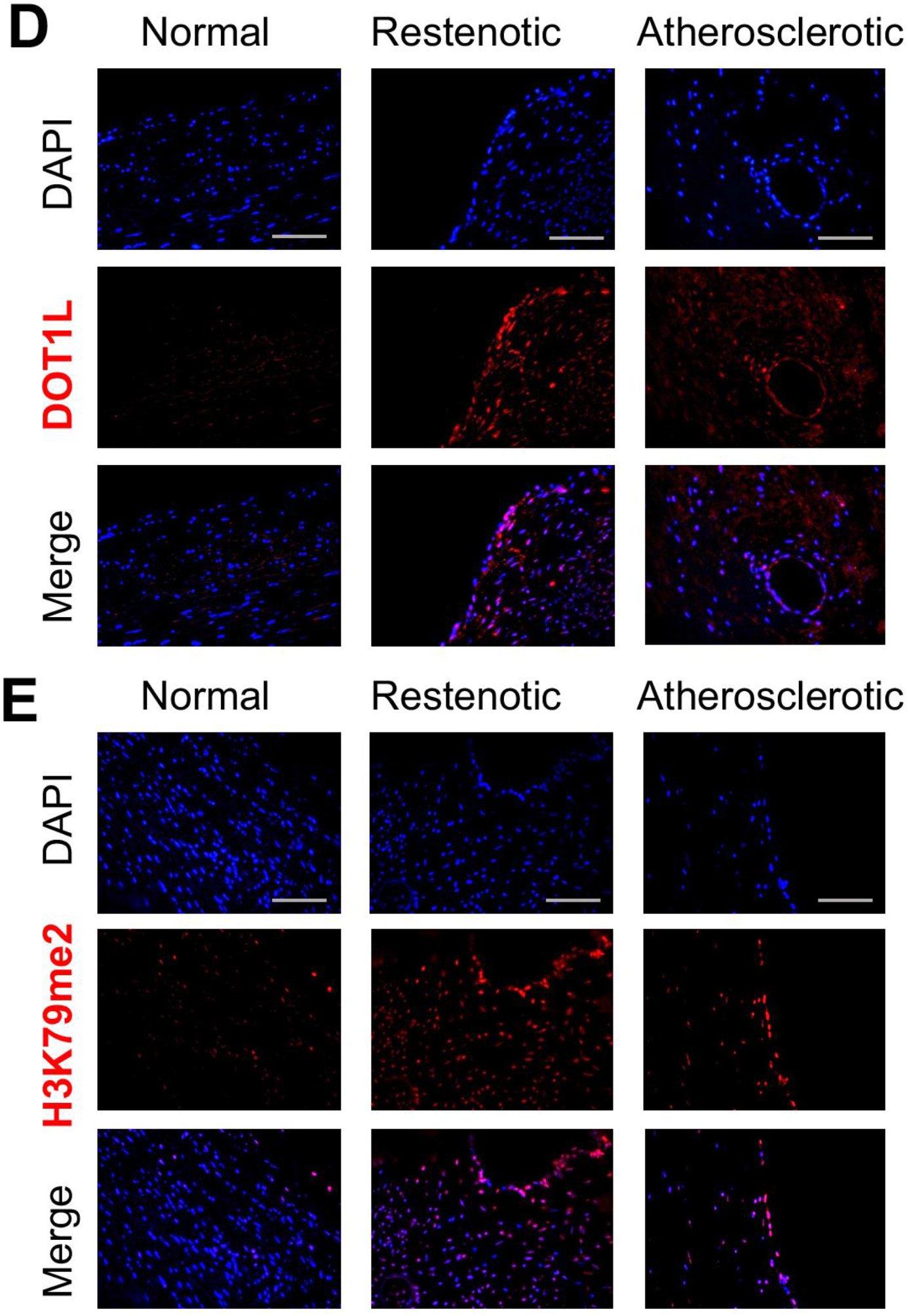
Upregulation of DOT1L and H3K79 methylation in balloon-injured arteries. A-C. Immunostaining of DOT1L, H3K79me2, and H3K79me3 on rat artery cross-sections. Balloon-injured rat common carotid arteries were harvested at post-injury days 3, 7, and 14. Uninjured contralateral arteries were used for control. D and E. Immunostaining on human artery samples (restenotic and atherosclerotic, normal artery for control). Negative staining without a primary antibody and distinguished rat artery wall morphological layers are shown in Figure S1. Scale bar: 100 μm; arrow points to the internal elastic lamina (IEL). A: adventitia; M: media; N: neointima. Quantification: Immunofluorescence intensity (measured by ImageJ above a threshold) was first normalized to the total number of DAPI-stained cells in each image field and then to the corresponding uninjured artery control. The data from different sections were pooled to generate the mean for each animal. The means from all animals in each group were then averaged, and the final mean ±SEM were calculated. Statistics: One-way ANOVA followed by Tukey’s post-hoc analysis; *P<0.05, **P<0.01, ***P<0.001; n (number of animals) is indicated by the data points in each group shown in the scatter plots; n.s., no significance; r.u., relative units.

Indicative of elevated DOT1L methyltransferase function after injury, staining of H3K79me2 and H3K79me3 exhibited temporal/spatial changes similar to that of the DOT1L protein (Figure 1, A-C). Importantly, increased staining of DOT1L and H3K79me2 was also observed in human restenotic and atherosclerotic (*vs* normal) artery tissues (Figure 1D and 1E; Figure S2) in which neointima typically develops^1^.

To determine a specific role for DOT1L in IH, we expressed shRNAs to silence DOT1L in the injured (endothelial-denuded) artery wall via luminal infusion with lentivirus. The transgene effectively reduced DOT1L protein and H3K79me2 (immunostaining) by 54.5% and 37.1%, respectively, as compared to scrambled shRNA control (Figure 2, A and B). This DOT1L silencing led to a 76.5% decrease of IH and nearly doubled lumen size (Figure 2C).

**Figure 2.**
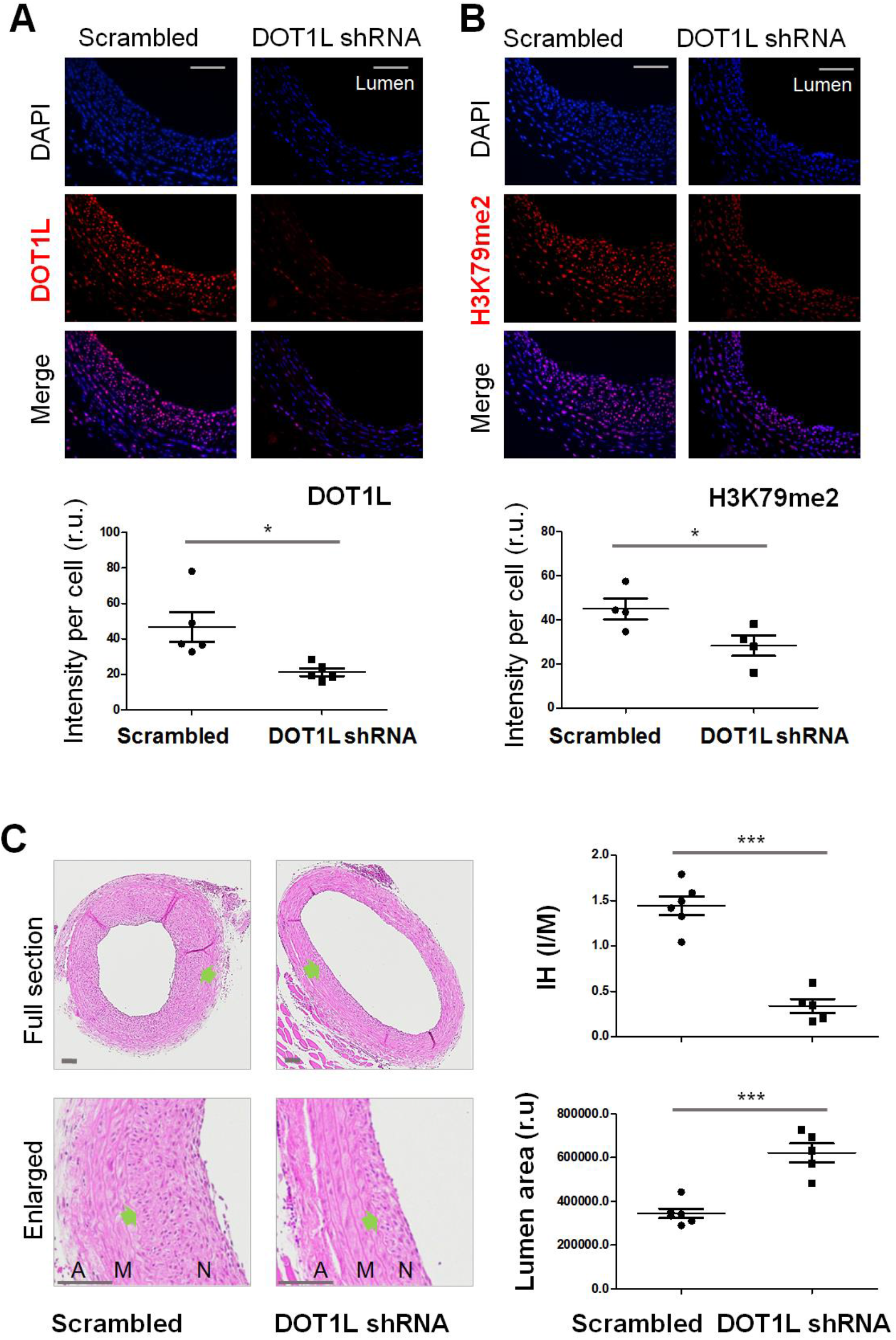
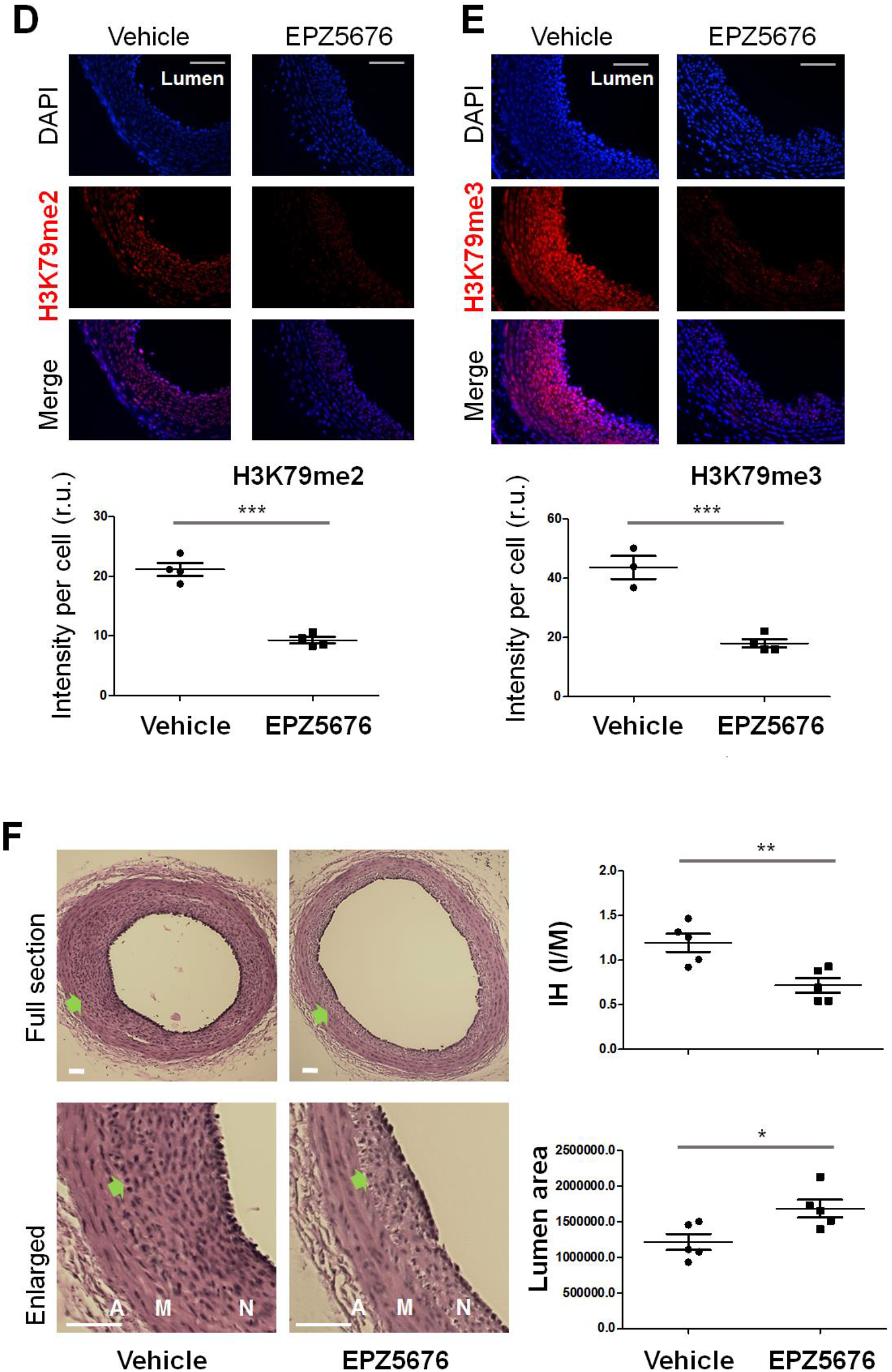
IH mitigation by DOT1L silencing or pharmacological inhibition. The IH model, DOT1L silencing or inhibition, and morphometric analysis of IH were performed as described in detail in Supplemental Materials. Immunostaining and data quantification to obtain mean ±SEM are described in Figure 1. A-C. In vivo DOT1L silencing via shRNA transgene. Immunostaining (and quantification) of DOT1L (A) and H3K79me2 (B) verifies effective DOT1L knockdown following DOT1L-specific shRNA transgene; morphometric analysis (C) indicates reduced IH in balloon-injured arteries infused with DOT1L-specific shRNA *vs* scrambled shRNA control. D-F. Periadventitial application of a novel DOT1L-selective inhibitor (EPZ5676, 37000-fold selectivity over other methyltransferases tested^7^). Immunostaining assay (D and E) and morphometric analysis show reduced H3K79me2/3 and IH, respectively, in injured arteries treated with EPZ5676 *vs* vehicle control. Scale bar: 100 μm; arrow points to IEL. A: Adventitia; M: media; N: neointima. Statistics: Unpaired Student t-test; *P<0.05, **P<0.01, ***P<0.001; n (number of animals) is indicated by the data points in each group shown in the scatter plots. Mann–Whitney non-parametric test was run in parallel: A, P=0.0079; B, P=0.0571; C, P=0.0043 (IH) or 0.0043 (lumen); D, P=0.0286; E, P=0.0571; F, P=0.0159 (IH) or 0.0317 (lumen).

To test a pharmacotherapy effect, we administered a DOT1L methyltransferase-selective inhibitor (EPZ5676^7^) around the injured artery. We found that both H3K79me2 and H3K79me3 were markedly reduced (by 56.1% and 58.6%, respectively) in EPZ5676-treated arteries compared to vehicle control (Figure 2, D and E). These changes were accompanied by a 39.9% decrease of IH and 38.6% increase of lumen size (Figure 2F).

## Discussion

To the best of our knowledge this is the first in vivo study addressing the importance of DOT1L in the development of IH. Our data show that DOT1L (protein and methylation function) markedly increased in rat common carotid arteries following balloon injury; either genetically silencing or pharmacologically inhibiting DOT1L effectively reduced IH.

Consistent with the histone-methylation function of DOT1L^4^, increased DOT1L, H3K79me2, and H3K79me3 in the injured artery wall were all confined in the nuclei, as illuminated by overlapped immunostaining and DAPI staining. Of note, as seen on day-7 and day-14 sections, the cells positively stained either for DOT1L, H3K79me2, or H3K79me3 localized mainly in the neointima layer, particularly in peri-luminal regions which are adjacent to various cytokine stimulants rich in the circulation. This is reminiscent of the distribution pattern of proliferative smooth muscle cells. In addition to the in vivo observations, treatment of rat primary smooth muscle cells with EPZ5676 in vitro reduced the cyclin D1 protein (Figure S3), a known pro-proliferation marker. Thus, future studies in the context of IH are warranted for understanding the epigenetic mechanism underlying the role for DOT1L in IH observed herein.

In summary, our results indicate that silencing DOT1L or inhibiting its methyltransferase function mitigates IH in the rat carotid artery angioplasty model. The cellular and molecular mechanisms underlying the DOT1L pathogenic role and its therapeutic targeting deserve further investigation.

## Abbreviations

ANOVA: Analysis of variance
DAPI: 4′,6-diamidino-2-phenylindole
DOT1L: Disruptor of telomeric silencing 1-like
H3K79: Histone 3 lysine 79
IH: (Neo)Intimal hyperplasia

## Acknowledgments

None

## Highlights

- DOT1L and its catalytic products H3K79me2 and H3K79me3 increased due to injury in rat carotid arteries after balloon angioplasty.
- Silencing DOT1L in vivo inhibited H3K79 methylation and diminished injury-induced intimal hyperplasia.
- Treatment with a DOT1L inhibitor reduced H3K79 methylation and intimal hyperplasia.

