## supplemental file for "Silencing epigenetic writer DOT1L attenuates intimal hyperplasia"

*Short title: Role of DOT1L in intimal hyperplasia*

[#] These authors contributed equally to this work.

* Corresponding author:

Lian-Wang Guo, Ph.D.

Department of Surgery and Department of Physiology & Cell Biology
The Ohio State University, Columbus, OH 43210, USA

The authors declare no conflict of interest

### Detailed Methods

***Animals.*** All animal studies conform to the *Guide for the Care and Use of Laboratory Animals* (National Institutes of Health) and protocols have been approved by the Institutional Animal Care and Use Committee at The Ohio State University (Columbus, Ohio). Male Sprague-Dawley rats purchased from Charles River Laboratories (Wilmington, MA) were used for experiments (at 300-350 g of body weight).

#### *Rat carotid artery balloon angioplasty*

The (neo)intimal hyperplasia (IH) model of rat carotid artery balloon angioplasty was performed as we previously described with minor modifications[1]. Briefly, rats were kept anesthetized (throughout the procedure) with 2-2.5% isoflurane via inhaling at a flow rate of 2 L per min. A neck skin incision was made and the left common carotid artery was dissected out. A 2-F balloon catheter (Edwards Lifesciences, Irvine, CA) was inserted into the common carotid artery through an arteriotomy on the left external carotid artery. The balloon was inflated to a pressure of 1.5 atm, withdrawn to the carotid bifurcation, and then deflated. This action was repeated four times, with the last withdrawal accompanied with balloon rotation (~3/4 round). These procedures cause injury to the artery (overstretch and endothelium denudation) that induces IH. DOT1L inhibition in injured arteries via gene silencing or a small molecule inhibitor was performed immediately following balloon angioplasty as described below. Blood flow was finally resumed and the neck incision was closed. The animal was kept on a 37°C warm pad to recover. For postoperative analgesia, in addition to carprofen and bupivacaine, buprenorphine (0.03 mg/kg) was subcutaneously injected.

#### *Lentiviral vector construction for DOT1L silencing*

To construct a lentiviral vector for the expression of DOT1L-specific shRNAs, the pLKO.1-puro empty vector was purchased from Addgene (Watertown, MA). A scrambled shRNA control and shRNAs specific for the rat DOT1L gene were designed by RNAi Central (http://cancan.cshl.edu/RNAi_central/step2.cgi). Three most efficient sequences (based on siRNAs as final products) are listed in Table S1. The corresponding shRNA-expressing lentivectors were constructed by using the pLKO.1-puro vector as a template. Lentiviruses were packaged in Lenti-X 293T cells (cat#632180, Clontech, Mountain View, CA) using a three-plasmid expression system (pLKO.1-shRNAs-puro, psPAX2 and pMD2.G) as described in our recent reports[1, 2], and used in combination (1:1:1) for carotid artery luminal infusion.

#### *DOT1L silencing in balloon injured carotid arteries*

In vivo DOT1L silencing was performed via luminal infusion of lentiviruses (into the denuded artery wall) to express DOT1L-specific shRNAs or a scrambled shRNA. We followed the infusion procedures described in our recent report[3]. Briefly, immediately following balloon angioplasty, a BD Insyte Autoguard IV catheter (24GA, BD, Franklin Lakes, NJ) was inserted into the injured segment of common carotid artery and suture-ligated together with the external carotid artery to prevent liquid leaking. The catheter was stabilized using a flexible magnetic mount arm platform (Quadhands, Charleston, SC). The lentivirus (>2.0 x $10^5$ IFU/ml) was injected with a syringe into the common carotid artery through the catheter, and the luminal infusion lasted for 25 min. The common carotid artery lumen was first flushed by temporally unclamping the proximal common carotid artery and internal carotid artery and then flushed with a 20 USP/ml heparin saline solution. The external carotid artery was permanently ligated and blood flow in both the common and internal carotid arteries was resumed. The surgery was finished as described above. To prevent thrombosis possibly caused by luminal infusion, 50 USP of heparin was subcutaneously administered prior to neck skin incision.

#### *Perivascular administration of a DOT1L-selective inhibitor to injured carotid arteries*

A selective DOT1L methyltransferase inhibitor (EPZ5676)[4] was periadventitially administered, as described in our previous report[1]. Briefly, balloon angioplasty was performed as described above except that the 4$^{th}$ balloon withdrawal was not included in this experiment which was performed at a different time than the lentivirus infusion experiment. After the surgery, the external carotid artery was permanently ligated and blood flow was resumed in the common and internal carotid arteries. EPZ5676 (10 mg/rat) or an equal amount of DMSO (vehicle control) was dispersed in mixed thermosensitive hydrogels (200 μl Triblock gel, AK12, Akina Inc., IN, and 200 μl Pluronic gel, Sigma, St. Louis, MO). The mix was then applied around the balloon-injured carotid artery. Surgery was finished as described above for the angioplasty model.

#### *Morphometric analysis of IH*

At post-injury day 3, 7, and 14, the injured left and uninjured contralateral common carotid arteries were excised from anesthetized animals following perfusion fixation at a physiological pressure of 100 mm Hg. Animal euthanasia immediately followed in a chamber gradually filled with $CO_2$. Cross-sections were prepared from paraffin-preserved common carotid arteries (5 μm thick each, excised from distal, middle, and proximal regions). The sections were used for immunostaining (see below) or Verhoeff-Van Gieson (VVG) or Haemotoxylin and Eosin (H&E) staining followed by morphometric analysis as we previously descibed[1]. Planimetric parameters as follows were measured on the sections. Lumen area; EEL area (inside external elastic lamina); IEL area (inside internal elastic lamina); (neo)intima area (IEL area minus lumen area); media area (EEL area minus IEL area). (Neo)intimal hyperplasia (IH) was quantified as a ratio of intima area versus media area (I/M). Measurements were performed using Image J by a student blinded to the experimental conditions. For each animal, 3–6 sections were used. The data from these sections were pooled to generate the mean for each animal. The means from all animals in each group were then averaged, and the standard error of the mean (SEM) was calculated.

#### *Immunostaining on artery tissue sections*

Fluorescent immunostaining was performed following our published protocol[3]. Briefly, artery sections were incubated without (negative staining) or with a primary antibody for 12 h and rinsed at least 3 times. The sections were then incubated with an anti-rabbit/mouse secondary antibody conjugated with Alexa Fluor 594 (A-11037/A-21203, Invitrogen, Carlsbad, CA) and rinsed. The specific antigen was then visualized with fluorescence microscopy. Detailed information of antibodies is included in Table S2. For quantification, 5 immunostained sections from each animal were used. Fluorescence intensity in each image field was quantified by using an ImageJ software and normalized to the number of DAPI–stained nuclei in the media and neointima layers. The values from all 5 sections were pooled to generate the mean for each animal. The means from all animals in each group were then averaged, and the final mean (±SEM) was calculated.

***Statistical Analysis.*** Data are presented as mean ± standard error of the mean (SEM). For statistical analysis, one-way ANOVA followed by post-hoc Tukey's test was applied for multi-group comparison and unpaired Student t-test (and also Mann–Whitney non-parametric test) was used for 2-group comparison, as specified in each figure legend. Statistical significance was set at $P < 0.05$.

**Supplemental figures**

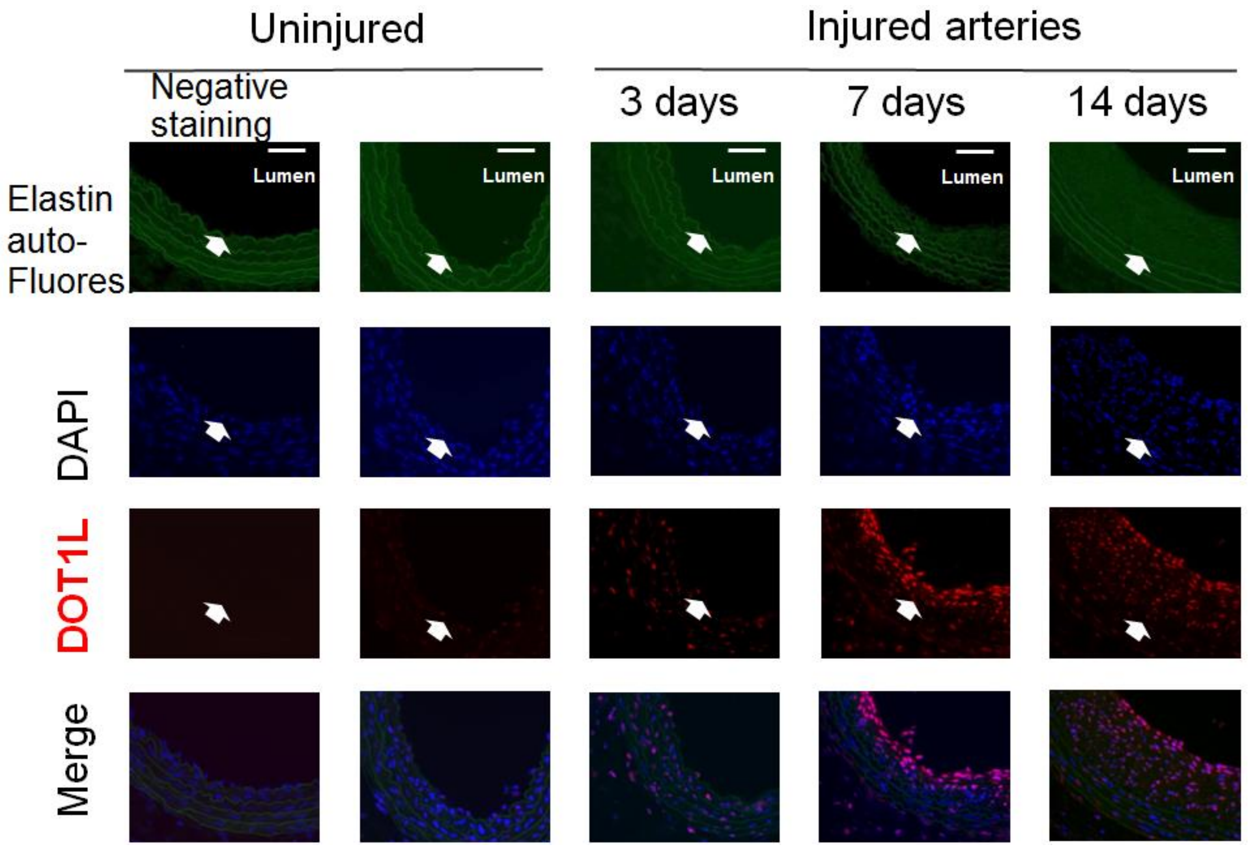

**Figure S1. Negative staining on rat carotid artery sections**

IH experiments and immunostaining were performed as described in Figure 1. Negative staining was performed with uninjured arteries without the use of a primary antibody. Both elastin autofluorescence and positive DOT1L staining were imaged to track the morphological layers of the artery. Arrow indicates the internal elastic lamina (IEL). The neointima layer is between the lumen and IEL. Scale bar: 100 $\mu$m. This figure correlates to Figure 1A.

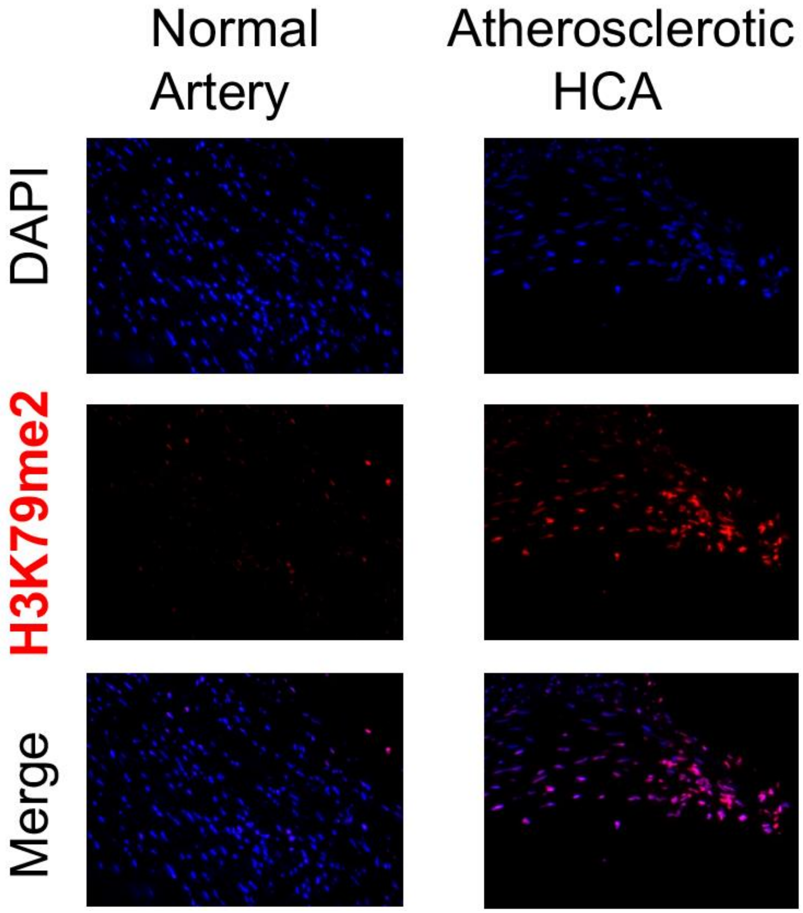

**Figure S2. Upregulation of H3K79me2 in human atherosclerotic coronary artery.**

Immunostaining was performed as described for Figure 1, D and E. Sections from a de-identified atherosclerotic coronary artery sample and a healthy control were subject to immunofluorescence staining for H3K79me2. (Please refer to human atherosclerotic femoral artery samples in Figure 1E).

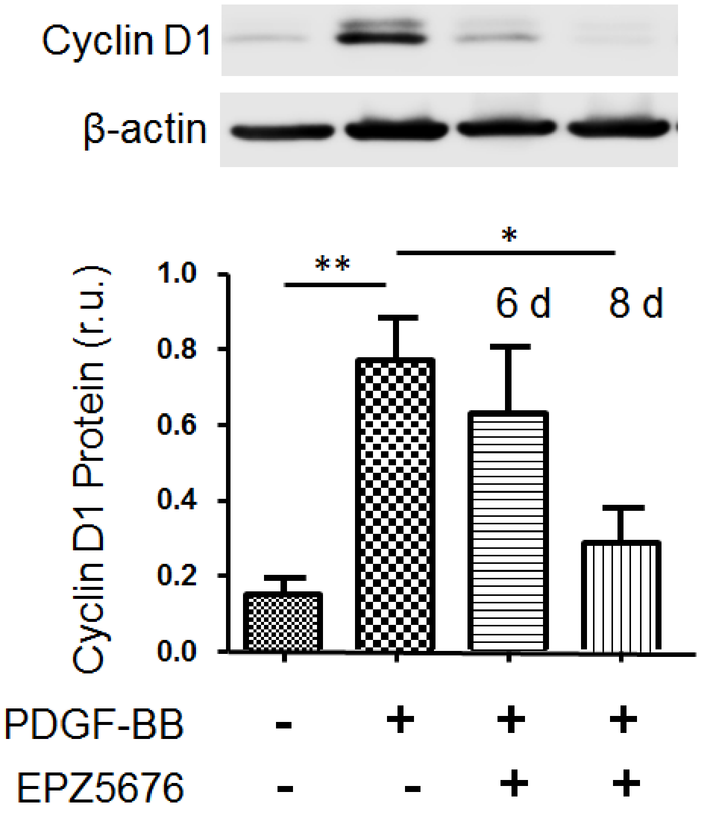

**Figure S3.** **Pretreatment with the DOT1L inhibitor EPZ5676 reduces PDGF-stimulated cyclin D1 expression**

Rat primary aortic artery smooth muscle cells were isolated, cultured, and used at passage-5 as we previously described[1]. To stimulate cell proliferation, the cells were incubated with platelet-derived growth factor (PDGF-BB, 20ng/ml) for 72h in the starvation medium (0.25% FBS) before harvesting for Western blot analysis. For pretreatment with DOT1L inhibitor, cells were cultured with EPZ5676 (10 μM) or DMSO vehicle control for 3 or 5 days in the starvation medium prior to adding PDGF-BB. Thus, cells were exposed to EPZ5676 for total 6 and 8 days before harvest. These two time-points were designed to match the post-injury day-7 peak time of DOT1L upregulation in the rat IH model. Detailed procedures of Western blot assay were described in our previous report[1]. Antibody sources are presented in Table S3.
Data quantification: densitometry was normalized to β-actin; one-way ANOVA followed by Tukey's post-hoc analysis; mean ±SEM; n =3 independent repeat experiments; *$P<0.05$, **$P<0.01$. This result is consistent with a slow turnover of H3K79 methylation (half-life ~ 3 days) in contrast to other histone modifications (e.g. acetylation turnover within a couple of hours)[5].

### Supplemental tables

**Table S1.** Scrambled and DOT1L-specific shRNAs (siRNA sequences)

| | |
|---|---|
| Scrambled shRNA Forward (F) | CCGGCCTAAGGTTAAGTCGCCCTCGCTCGAGCGAGGGCG ACTTAACCTTAGGTTTTTG |
| Scrambled shRNA Reverse (R) | AATTCAAAAACCTAAGGTTAAGTCGCCCTCGCTCGAGCGA GGGCGACTTAACCTTAGG |
| rDOT1L shRNA1F | CCGGACCAGAGAAGCTCAACAACTACCTCGAGGTAGTTGT TGAGCTTCTCTGGGTTTTTG |
| rDOT1L shRNA1R | AATTCAAAAAACCAGAGAAGCTCAACAACTACCTCGAGGTA GTTGTTGAGCTTCTCTGGG |
| rDOT1L shRNA2F | CCGGCGCTGTTAGAGTCCTTCAAGATCTCGAGATCTTGAA GGACTCTAACAGCTTTTTTG |
| rDOT1L shRNA2R | AATTCAAAAACGCTGTTAGAGTCCTTCAAGATCTCGAGATC TTGAAGGACTCTAACAGCT |
| rDOT1L shRNA3F | CCGGACCAGAGCATGCCAAGGAGAACCTCGAGGTTCTCCT TGGCATGCTCTGGCTTTTTG |
| rDOT1L shRNA3R | AATTCAAAAAACCAGAGCATGCCAAGGAGAACCTCGAGGT TCTCCTTGGCATGCTCTGGC |

**Table S2**. Antibodies used for immunostaining

| Antigen | company | Catalog number | Dilution ratio |
|---|---|---|---|
| DOT1L | Cell Signaling Technology | 90878s | 1:100 |
| DOT1L | Abnova | H00084444-A01 | 1:100 |
| H3K79me2 | Cell Signaling Technology | 9847T | 1:100 |
| H3K79me3 | Novus Biologicals | NB211383SS | 1:100 |

**Table S3**. Antibodies used for Western blotting

| Antigen | company | Catalog number | Dilution ratio |
|---|---|---|---|
| cyclinD1 | Abcam | Ab134175 | 1:10000 |
| β-actin | Abcam | Ab6276 | 1:3000 |
